# netDx: Interpretable patient classification using integrated patient similarity networks

**DOI:** 10.1101/084418

**Authors:** Shraddha Pai, Shirley Hui, Ruth Isserlin, Muhammad A Shah, Hussam Kaka, Gary D Bader

## Abstract

Patient classification has widespread biomedical and clinical applications, including diagnosis, prognosis and treatment response prediction. A clinically useful prediction algorithm should be accurate, generalizable, be able to integrate diverse data types, and handle sparse data. A clinical predictor based on genomic data needs to be easily interpretable to drive hypothesis-driven research into new treatments. We describe netDx, a novel supervised patient classification framework based on patient similarity networks. netDx meets the above criteria and particularly excels at data integration and model interpretability. As a machine learning method, netDx demonstrates consistently excellent performance in a cancer survival benchmark across four cancer types by integrating up to six genomic and clinical data types. In these tests, netDx has significantly higher average performance than most other machine-learning approaches across most cancer types and its best model outperforms all other methods for two cancer types. In comparison to traditional machine learning-based patient classifiers, netDx results are more interpretable, visualizing the decision boundary in the context of patient similarity space. When patient similarity is defined by pathway-level gene expression, netDx identifies biological pathways important for outcome prediction, as demonstrated in diverse data sets of breast cancer and asthma. Thus, netDx can serve both as a patient classifier and as a tool for discovery of biological features characteristic of disease. We provide a software complete implementation of netDx along with sample files and automation workflows in R.

## Introduction

The goal of precision medicine is to build quantitative models that guide clinical decision-making by predicting disease risk and response to treatment using data measured for an individual. Within the next five years, several countries will have general-purpose cohort databases with 10,000 to >1 million patients, with linked genetics, electronic health records, metabolite status, and detailed clinical phenotyping; examples of projects underway include the UK BioBank^1^, the US NIH Precision Medicine Initiative (www.whitehouse.gov/precision-medicine), and the Million Veteran Program (http://www.research.va.gov/MVP/). Additionally, human disease specific research projects are profiling multiple data types across thousands of individuals, including genetic and genomic assays, brain imaging, behavioural testing and clinical history from integrated electronic medical records^2–4^ (e.g. the Cancer Genome Atlas, http://cancergenome.nih.gov/). Computational methods to integrate these diverse patient data for analysis and prediction will aid understanding of disease architecture and promise to provide actionable clinical guidance.

Statistical models that predict disease risk or outcome are in routine clinical use in fields such as cardiology, metabolic disorders, and oncology^5–8^. Traditional clinical risk prediction models typically use generalized linear regression or survival analysis, in which individual measures are incorporated as terms (or features) of a single equation. Standard methods of this type have limitations analyzing large data from genomic assays (e.g. whole-genome sequencing). Machine learning methods can handle large data, but are often treated as black boxes that require substantial effort to interpret how specific features contribute to prediction. Black box methods are unlikely to be clinically successful, as physicians frequently must understand the characteristic features of a disease to make a confident diagnosis^9^. Interpretability is particularly required in genomics because of relatively smaller sample sizes and to better understand the molecular causes of disease. Further, many existing methods do not natively handle missing data, requiring data pruning or imputation, and have difficulty integrating multiple different data types.

The patient similarity network framework can overcome these challenges and excels at integrating heterogeneous data and generating intuitive, interpretable models. In this framework, each feature of patient data (e.g. gene expression profile, age) is represented as a patient similarity network (PSN) (Figure 1A). Each PSN node is an individual patient and an edge between two patients corresponds to pairwise similarity for a given feature. For instance, two patients could be similar in age, mutation status or transcriptome. PSNs can be constructed based on any available data, using a similarity measure (e.g. Pearson correlation, Jaccard index). Because all data is converted to a single type of input (similarity networks), integration across diverse data types is straightforward. Patient similarity networks (PSN) are a recently introduced concept and have been used successfully for unsupervised class discovery in cancer and type 2 diabetes^10,11^, but have never been developed for supervised patient classification.

**Figure 1.**
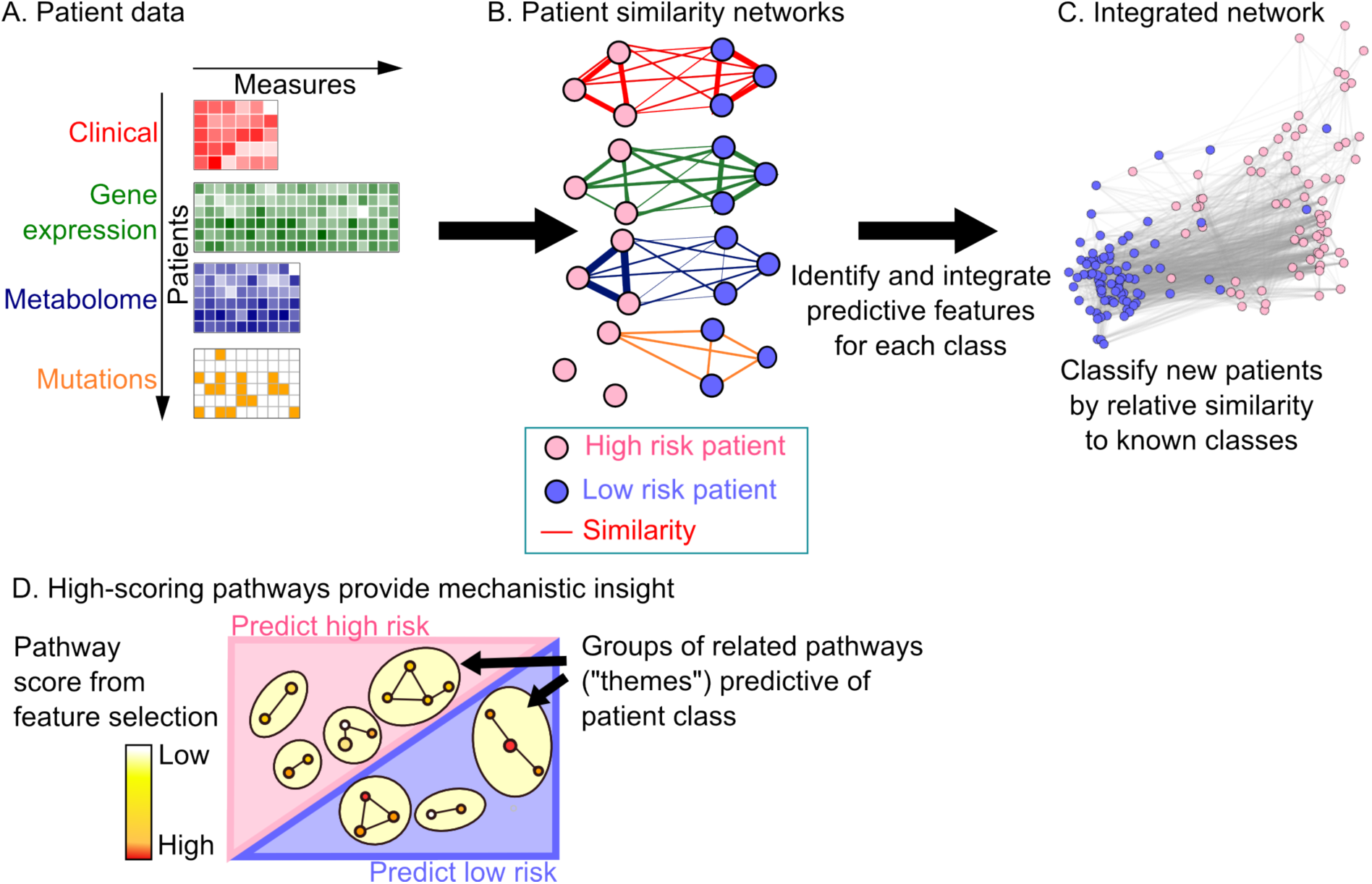
The netDx method. netDx converts patient data (A) into a set of patient similarity networks (PSN), with patients as nodes and weighted edges representing patient similarity by some measure (B). The simple example for predicting low/high risk for disease uses clinical, genomic, metabolomic and genetic data. netDx identifies which networks strongly relate high-risk patients (here, clinical and metabolomic data) and which relate low-risk patients (clinical and gene expression data). Feature selection is used to score networks by their ability to predict patient class; details in Figure S1. C. netDx returns several types of output. Top-scoring features are combined into a single view of overall patient similarity, which can be used to classify new patients based on relative similarity to known patient classes. netDx also provides standard classifier performance metrics and scores for the predictive value of individual features. D. If pathway features are used, netDx identifies and visualizes the pathways most useful for classification.

We describe netDx, the first PSN-based approach for supervised patient classification. In this system, patients of unknown status can be classified based on their similarity to patients with known status. This process is clinically intuitive because it is analogous to clinical diagnosis, which often involves a physician relating a patient to a mental database of similar patients they have seen. As demonstrated below, netDx has strengths in classification performance, heterogeneous data integration, usability and interpretability.

## Results

### Algorithm Description

The overall netDx workflow is shown in Figure 1. This example conceptually shows how PSNs can be used to predict if a patient is at high or low risk of developing a disease based on a variety of patient-level data types. Similarity networks are computed for each patient pair and for each data type. In this example, high-risk patients are more strongly connected based on their clinical profile, which may capture age and smoking status, and metabolomics profile. Low-risk patients are more similar in their clinical and genomic profiles. The goal of netDx is to identify the input features predictive of high and low risk, and to accurately assign new patients to the correct class.

#### Input data design

Each patient similarity network (PSN) is a feature, similar to a variable in a regression model (we use the terms “input networks” and “features” interchangeably). A PSN can be generated from any kind of patient data, using a pairwise patient similarity measure (Figure 1A). For example, gene expression profile similarity can be measured using Pearson correlation, while patient age similarity can be measured by the normalized difference. A reasonable design is to define one similarity network per data type, such as a single network based on correlating the expression of all genes in the human genome, or a network based on similarity of responses to a clinical questionnaire. If a data type is multivariate, defining a network for each individual variable will result in more interpretable output. However, this approach may lead to too many features generated (e.g. millions of SNPs), which increases computational resource requirements and risk of overfitting. Thus, as with any machine learning task, there is a trade-off between interpretability and overfitting/scalability. To help address this problem for gene-oriented data (e.g. transcriptomics), we group gene-based measurements into biological pathways, which we assume capture relevant aspects of cellular and physiological processes underlying disease and normal phenotypes. This biological process-based design generates ∼2,000 networks from gene expression profiles containing over 20,000 genes, with one network per pathway.

#### Selecting features informative of class prediction

Feature selection identifies the input networks with the highest generalizable predictive power, and is run once per patient class. netDx is trained on samples from the class of interest, using cross-validation (Figure S1) and an established association network integration algorithm^12,13^. The algorithm scores each network based on its value in the classification task. The ideal network is one connecting all patients of the same class without any connections to other classes; for example, one connecting all treatment responders, and all non-responders, without edges between the two. The least useful network is one that connects patients from one class to patients from other classes, without connecting any patients in the same class; for example, one that connects responders and non-responders to the same extent. In each cross-validation fold, regularized linear regression assigns network weights, reflecting the ability to discriminate query patients from others, and removes uninformative networks. netDx increases a network’s score based on the frequency with which it is assigned a positive weight in multiple cross-validation folds. The classifier’s sensitivity and specificity can be tuned by thresholding this score; a network with a higher score achieves greater specificity and lower sensitivity. The output of this feature selection step is a set of networks that can be integrated to produce a predictor for the patient class of interest.

#### Class prediction using selected features

After training and feature selection are separately run for each class, feature selected networks are combined by averaging their similarity scores to produce an integrated network. Test patients are ranked by similarity to each class using label propagation in the integrated network, and are assigned to the class with the highest rank^14,15^ (Figure S2).

#### netDx output (Figure 1C-D)

netDx returns predicted classes for all test patients and standard performance measures including the area under the receiver operating characteristic curve (AUROC), area under the precision-recall curve (AUPR), and accuracy. Scores for each feature are returned and if pathway features are used, they are visualized using an enrichment map (Figure 1D)^16^. The integrated patient network is visualized and used to assess the strength of class separation, and inter-and intra-class separation is measured using average shortest path methods (Online Methods, Figure 1C).

#### Benchmarking performance by predicting binarized survival in cancer

To assess the classification performance of netDx, we use an established cancer survival prediction benchmark available for four tumor types, using data from The Cancer Genome Atlas (TCGA; http://cancergenome.nih.gov/) via the TCGA PanCancer Survival Prediction project website of Yuan et al.^17^, https://www.synapse.org/#!Synapse:syn1710282, Table 1). These tumor types have been thoroughly analyzed using eight machine learning methods, which provides extensive performance results that we can compare to^17^. Data are for renal clear cell carcinoma^18^ (KIRC, N=150 patients), ovarian serous cystadenocarcinoma^19^ (OV, N=252), glioblastoma multiforme^20^ (GBM, N=155), and lung squamous cell carcinoma^21^ (LUSC, N=77). Data for a given tumour type includes: clinical variables (e.g. age, tumour grade); mRNA, miRNA and protein expression; DNA methylation; and somatic copy number aberrations. Binarization of survival and format of clinical variables followed previous work^17^.

For each tumour type, we classified high and low survival using multiple combinations of input data following the original work^17^. For each model tested, patient samples were split 80:20 into a training and a test set. Using only the training samples, 10-fold cross validation was performed for each class (good survival; poor survival), generating for each network a score between 0 and 10. The best scoring networks (9 or 10) were selected as features and used to classify test samples. This process was repeated for random splits of train and test until performance measures stabilized (20 splits) (Figure 2A). Predictor performance was measured as the average of test classification across the 20 splits. We tested fifteen different predictor models that varied by prefiltering strategy, choice of similarity metric, and whether networks were defined at the level of individual variables (genes) or entire datatypes (Supplementary Table 1). Prefiltering is an initial feature selection step which creates patient similarity networks only using variables that pass lasso regression; it is performed within the cross-validation loop to avoid leaking information between train and test samples. For similarity metrics, we tested normalized difference, Pearson correlation with and without exponential scaling, radial basis function, and Euclidean-distance based similarity with exponential scaling (see Online Methods).

**Figure 2.**
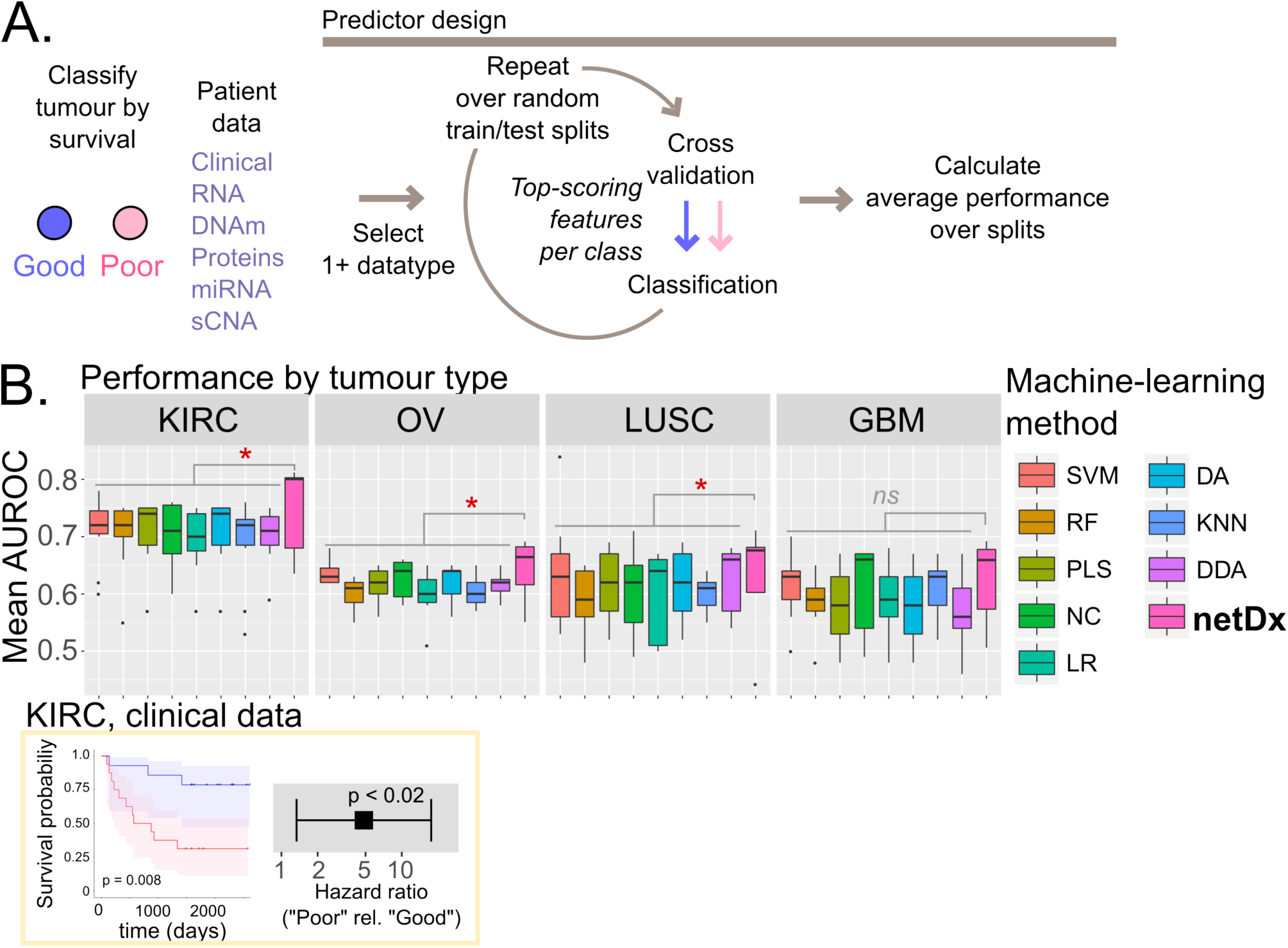
Performance benchmarking using the PanCancer Survival data A. Method. Various combinations of networks were input to netDx to predict binary survival (YES/NO). Predictors were built using 20 random train/test splits. B. Average performance of netDx compared to other machine-learning methods. Each panel shows data for one tumour type, and a boxplot shows mean AUROC for a given machine-learning method for different tested combinations of input data (Supplementary Table 1). netDx is shown in pink. As a reference point, Kaplan-Meier curves and hazard ratios are shown for predicted samples from a representative KIRC split (bottom). P-values are from one-sided Wilcoxon-Mann-Whitney comparing netDx to all other methods combined.

We use the AUROC to measure performance, as this was the metric reported by the PanCancer survival project^17^. Information on the exact samples used for the 10 train/test splits used by Yuan et al. is not available, thus we used new random splits, but used more splits (n=20) chosen to reach a stable AUROC estimate. netDx demonstrates consistently excellent performance for all four tumour types (Figure 2B); average netDx performance is significantly higher than that for all other methods for three of the tumours (one-tailed WMW; KIRC: p < 0.03; OV: p < 0.013; LUSC: p < 0.04), and is close to significant for the fourth (GBM: p < 0.06). Further, the top-performing netDx model outperforms all eight tested machine learning algorithms for kidney and ovarian cancer (Figure 2B, Table S1), performs at par for brain cancer (netDx best=0.69; Yuan et al. best=0.71), and outperforms all but one outlier data point for lung cancer. Performance statistics reported for Yuan et al. were the best performing models out of hundreds tested for different data combinations with eight different machine learning methods: diagonal discriminant analysis; K-nearest neighbor; discriminant analysis; logistic regression; nearest centroid; partial least squares; random forest; and support vector machine. Thus, netDx performs as good or better than a diverse panel of machine-learning methods and parameter combinations.

### Pathway-level feature selection identifies cellular processes predictive of clinical condition

Creating a single feature per datatype identifies the general predictive value of that data layer but, without further work, does not provide insight into which genes or cellular processes are useful for classification. This information is useful to better understand the mechanisms of disease. netDx natively supports the ability to group unit measures into relevant groupings for more interpretable features. For instance, genes can be grouped into pathways (gene sets) so that the feature selection process scores pathways for predictive value. To illustrate this ability, we classified breast tumours as being of the Luminal A subtype or not, starting from tumour-derived gene expression (N=348 patients; (154 Luminal A, 194 non-Luminal A^22^). Gene expression was used as input and features were defined at the level of curated pathways of cellular processes. Cross-validation was run with 100 80:20 train/test splits. Each split included a feature selection process that resamples the training data ten times; each split therefore scores features between 0 and 10, with higher values indicating greater predictive power. We then calculated the highest score each feature consistently obtained over all 100 splits (here defined as the highest score obtained for >=70% of the 100 splits) as a stable measure of that feature’s predictive power. Resulting pathways were visualized as an EnrichmentMap (Figure 3).

**Figure 3.**
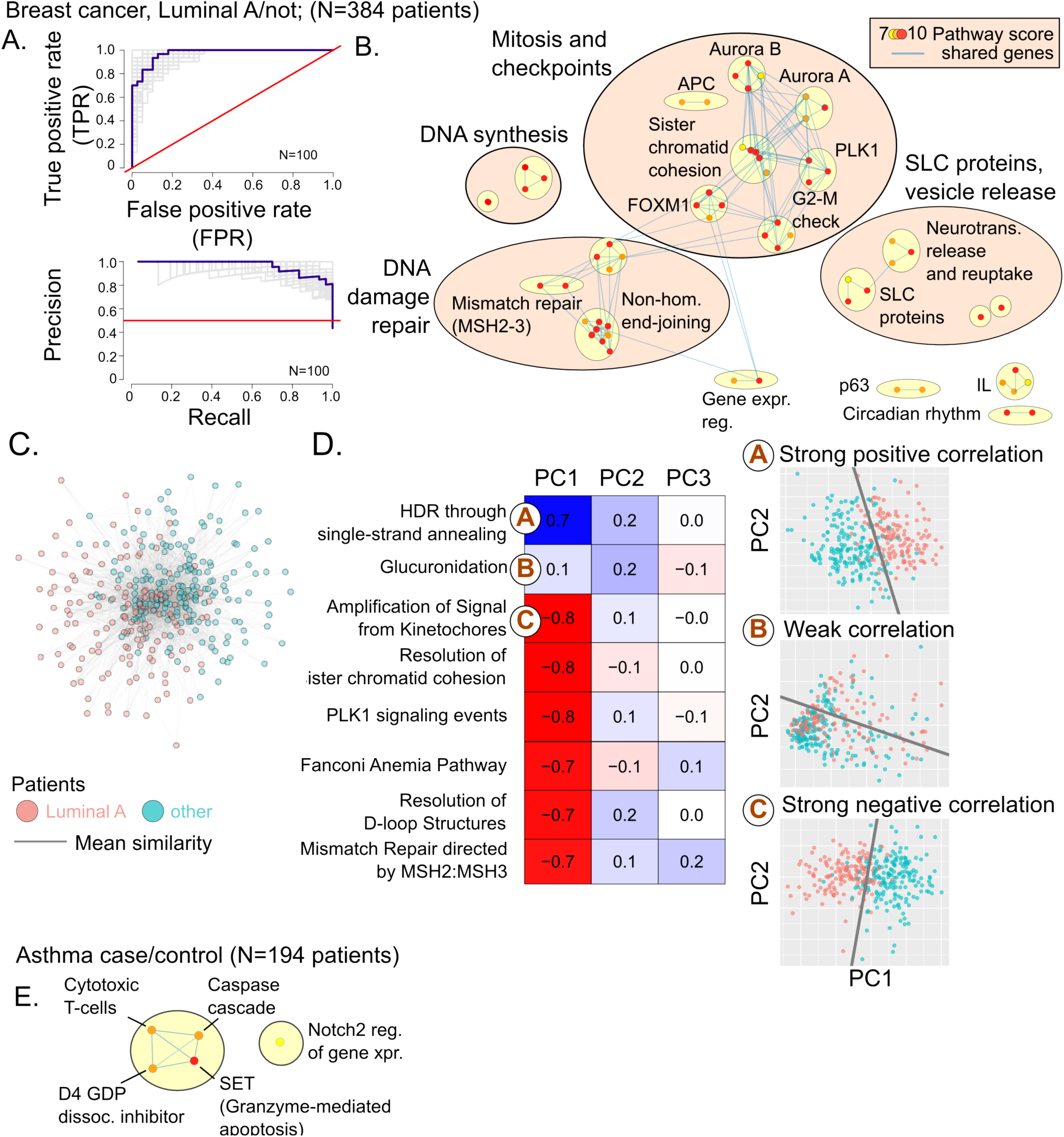
Pathway-level feature selection in breast cancer and asthma A. netDx Performance for binary classification of breast tumour as Luminal A subtype from tumour-derived gene expression (N=384 patients). B. Pathways feature-selected by netDx in predicting Luminal A status. Nodes are pathways and edges indicate shared genes. Nodes are coloured by highest netDx score consistently achieved out of a maximum possible of 10, in >=70% of 100 train/test splits. Themes identified by AutoAnnotate^16,37^. C. Integrated patient similarity network. Nodes are patients, and edges are average similarity from the pathways that scored 10 out of 10 in all splits. Nodes are coloured by tumour type. Edges with weight < 0.7 were excluded and the top 20% of edges per node were retained. The resulting network was visualized in Cytoscape (spring-embedded layout). D. Correlation of top-scoring pathway features (represented as the first three principal components of pathway-specific gene expression) with tumour type (Spearman’s correlation). Table cells are colored by sign and magnitude of correlation (blue: Spearman corr. **>**0; red, corr. <0). Circled letters correspond to detailed panels on the right. Right: Projections of patient-level gene expression in feature-selected pathways onto first two principal components (individual dots indicate patients). Points are colored by survival class. Decision boundaries were calculated using logistic regression on scatterplot data. E. Selected features for asthma case status in the case of asthma case/control prediction (N=97 cases; N=97 controls). Legend as in (B).

Tumour classification was near perfect, with an average (SD) AUROC of 0.97+/−0.01, average AUPR of 0.93+/−0.02, and average accuracy of 89% +/−3% (Figure 3A). Performance for pathway-level features is slightly, but significantly, better than when gene expression is provided as a single feature (mean AUROC for single network = 0.96+/−0.02; two-sided Wilcoxon-Mann-Whitney test p < 0.025). Top-scoring pathways included cell cycle progression and checkpoint regulation, DNA synthesis, DNA mismatch repair, and DNA double-strand break repair themes (Figure 3B, Table S2). These processes are consistent with the pathways known to be dysregulated in luminal breast tumours and cancer progression in general. netDx also identified pathways related to Solute Carrier Family membrane transport proteins and vesicle release, which are not traditionally linked to breast cancer, but may support new insights (see discussion). We integrated patient similarity networks from top-scoring pathways (those scoring 10 out of 10 in all splits) into a single patient similarity network (Figure 3C). In this network, LumA patients are significantly closer to other LumA patients (average shortest distance=0.52), compared to patients of other breast cancer subtypes (average shortest distance=0.58; one-tailed Wilcoxon-Mann-Whitney test p < 2e-16).

A common problem in clinical genomics is relating higher-level analyses, such as the set of affected pathways in Figure 3B, back to changes in individual patients. To address this problem, we performed principal component analysis on gene expression values of all genes within a given pathway and correlated the projections of the first three principal components with clinical outcome (Figure 3). Most features individually showed significant correlation with tumour subtype (e.g. correlation for “Amplification of Signal from the Kinetochores”=−0.80, p < 3.3e-72), and the patient class boundary is visually evident in these features (Figure 3D). However, not all features had this property (e.g. correlation for “Glucuronidation” = 0.1, p < 0.038). Pathways that score highly in feature selection and correlate with outcome are good candidates for follow-up biomarker or mechanistic studies.

As a second case study to demonstrate that netDx feature selection identifies pathways consistent with the biology of the condition, we predicted case/control status in asthma using gene expression from sorted peripheral blood mononuclear cells^23^ (97 cases, 97 controls) and an identical predictor design as used for breast cancer above. The netDx predictor achieved an AUROC of 0.71+/−0.07 (SD) (Figure 3E; mean AUPR=0.65; mean accuracy=66%). Selected pathways included cytotoxic T-lymphocytes related processes and Notch2 signaling (Figure 3D; Table S3). The feature-selected themes in breast cancer and asthma are each consistent with prior knowledge of cellular changes in these conditions (see discussion). These examples demonstrate that when used with pathway-level features, netDx can provide insight into the molecular mechanisms that discriminate between patient groups and into general disease related processes. Altogether, our results show that using pathway-level features can improve classification performance and provide insight into disease mechanisms.

## Discussion

We describe netDx, the first supervised patient classification system based on patient similarity networks. We demonstrate that netDx has excellent classification performance predicting survival across four different tumour types. Further, feature selection, especially when biological pathways are used, aids interpretability and provides insight into disease mechanisms important for classification. This framework can be used to create accurate, generalizable predictors, and has particular strengths in data integration and interpretation compared to other machine learning approaches. netDx is targeted at researchers who are interested to see if their patient-level data can answer a specific patient classification question. netDx provides a standard workflow that can quickly determine if the classification question can be answered based on a training set and if so, provides a set of relevant features and a software tool to classify new patients.

netDx demonstrated consistently excellent performance with the PanCancer benchmark set, performing as good or better in almost all cases. A single support vector machine model from Yuan et al. for lung cancer survival prediction vastly outperformed any other model, including netDx. This may be because the latter model identified a non-linear decision boundary or was overfit. Future versions of netDx will explore if considering non-linear effects (e.g. via non-linear similarity measure and network combinations) can improve performance.

netDx also includes support for feature grouping to improve interpretability while keeping feature number low. Reducing the number of features can mitigate overfitting risk, improve signal detection with sparse data, require less compute resources, and improve prediction performance. While we demonstrate this functionality by grouping gene-level expression measures into pathway-level features (Figure 3), any feature grouping is possible. Groupings with clear clinical or mechanistic interpretation will aid classification interpretation. The themes identified for Luminal A classification of breast tumours are consistent with processes known to be dysregulated in this type of cancer. For instance, themes of DNA repair and G2-M checkpoint regulation are consistent with the known roles of BRCA1/BRCA2 and ATM proteins, which are established risk factors for breast cancer^24^. Cell cycle dysregulation accompanies genomic instability as a feature of several cancers^25^. netDx also identified a theme of solute carrier family proteins, many of which are overexpressed in tumours, are thought to mediate the altered metabolic needs of growing tumours^26,27^, and are associated with genetic risk for breast cancer^28^. Even genes typically thought to be involved in neurotransmitter release are expressed in multiple TCGA cancers^27^; for instance, ABAT, which is responsible for catabolism of the neurotransmitter GABA, is a biomarker for poor hormone therapy outcome in advanced stages of breast cancer^29^. Therefore, pathways identified by netDx may suggest novel directions for biomarker identification and therapeutic targeting. Similarly, the themes identified for asthma case prediction, including cytotoxic T-lymphocytes and associated apoptosis, are consistent with known asthma genetics and genomics results. Asthma is an inflammatory condition affecting the airways of the lung. Genetic loci associated with asthma include genes involved in immunoregulation and T-helper cell differentiation, and T-cell activation has been identified in blood of affected individuals from transcriptomic and DNA methylation studies^23,30–32^. Notch signalling regulates the differentiation of T-helper cells, and inhibitors of this pathway are being tested in clinical trials to suppress symptoms of asthma^33–35^. In summary, when provided with pathway-level features, netDx can be a useful tool for discovery research.

Good performance and interpretability increase confidence of prediction results. However, it is sometimes difficult to know how well a prediction method is performing if there is nothing to compare with. Thus, we recommend that machine-learning classifiers, such as netDx, be assessed using a predictor checklist of tests to gain confidence in the classification results (Supplementary Figure 3). Such a checklist would include:

1. traditional performance metrics, including the AUROC, AUPR, F1, and accuracy
2. the extent to which the predictor captures prior knowledge about the disease under study, such as known cellular pathways
3. an orthogonal measure of the validity of the predicted classes. For instance, in the context of survival prediction in cancer, a predictor should result in significantly separable survival curves for the two predicted patient sets
4. a measure of the strength of separation of the classes, such as the extent to which patient classes separately cluster in the integrated similarity network
5. a measure of how much the results are better than random, measured using an appropriate set of negative controls

A list of passing or failing grades on each test would provide a simple report card to compare several predictor designs (Figure S4) and intuitive visualizations of the results (e.g. Figures 2B and 3) would improve understanding about how well the predictor may work in general.

netDx provides a complete framework for precision medicine. However the ultimate vision is to enable clinical researchers to assess classification performance for questions of interest, such as ‘will a patient respond to one therapy or another?’ based on patient measurements and outcomes present in large electronic medical record databases. Output would include a report card on model performance and generalizability estimates on independent cohorts, feature interpretation, an interactive integrated patient similarity network visualization enabling the exploration of individual patients, and a ready-to-run classifier for new patients.

netDx is implemented as an open-source R software package available at http://netdx.org, with worked examples. We also propose that users store and publicly share patient similarity networks, useful as features for netDx and other PSN methods, in the NDEx network exchange system^36^.

## Acknowledgements

We thank Quaid Morris and Daniele Merico for discussions on method development, and Lincoln Stein for feedback on the manuscript. We also thank Han Liang for discussion on implementation details for the machine learning used in Yuan et al. (2014). This work was supported by a Canadian Institutes of Health Research award to GDB (grant number 499509), by the NRNB (U.S. National Institutes of Health grant number 499382), by a grant for Cytoscape (U.S. National Institutes of Health grant number 503758), and by a Canadian Institutes of Health Research Fellowship award to SP (award number 498002).

## Author contributions

All authors contributed to netDx method development. S.P. and M.A.S. analyzed the data. S.P. wrote the netDx software package with contributions from M.A.S. S.H., H.K. and R.I. developed initial versions of netDx. S.P. and G.D.B. wrote the paper. G.D.B. supervised the work.

## Online Methods

### PanCancer Survival benchmark models

We tested various models for the PanCancer survival benchmarking exercise. This section describes the model details; models are named as per Supplementary Table 1. The models varied based on whether or not they included a data imputation step, whether or not variables were prefiltered using lasso regression, and choice of similarity metric (Pearson correlation, normalized difference, scaled Euclidean/Pearson). Where used, imputation was performed separately for training and test samples to avoid information leaking from train to test. Where used, prefiltering was performed on training samples within a cross-validation loop to avoid information leaking from train to test.

#### Base (no lasso prefiltering)

In this model, each datatype was treated as a single feature; i.e. one patient similarity network was generated for gene expression, one for clinical data, etc. Similarity was defined by Pearson correlation where a datatype had more than six measures^1^, or by average normalized difference if the datatype had five or fewer variables. For a set of *k* variables G={g_1_,g_2_,..g_k_}, where 1<=k<=5, the similarity *S* between two patients *a* and *b* is defined as the average of normalized differences for each of the variables:

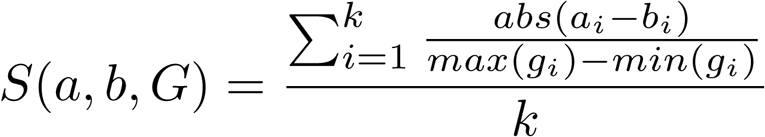

For the case of a single continuous variable, similarity is computed as normalized difference, defined as:

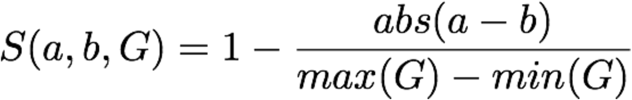

where *a* and *b* are the values of the variable for individual patients (a and b) and *G* is the set of all values for the variable (e.g. age).

#### Variable prefiltering and Scaled Euclidean / Scaled Pearson

This design combines within-CV prefiltering with lasso regression^2^, and defines features at the level of individual variables (e.g. genes, clinical variables). It enables netDx to score individually predictive variables in contrast to combining all variables of a data type into a single network, and is likely a better choice when signal is not widespread in a datatype. Within each cross-validation fold, lasso regression was applied to training samples for each datatype (prefiltering), and only variables with a non-zero weight were included. Regression used only training samples within a given fold to avoid leaking information from test to train. The similarity metric used is either Euclidean distance (model code= *euc6K*) or Pearson correlation, followed by local exponential scaling ^3^. Imputing missing data by median further improved performance only for glioblastoma (*eucimpute, pearimpute*). Imputation was performed within cross-validation, and was performed separately for training and test samples to avoid leaking information from train to test. The lung cancer dataset demonstrated the best performance if the model was also limited to the top clinical variable from lasso (*plassoc1*).

### Integrated patient network

The integrated patient network is an average combination of all feature-selected networks to create a single network (i.e. average of all edge weights between patients from all selected networks). Visually, the goal is to view more similar patients as being more tightly grouped, and more dissimilar patients as being farther apart. Similarity is therefore converted to dissimilarity, defined as 1-similarity. Weighted shortest path distances are computed on this resulting dissimilarity network. To aid visualization, only edges representing the top 20% of distances in the network are included. For the network with a single clinical network, the top 50% of distances are included, to limit the number of patients without edges.

### Survival curve and hazard ratios

Survival curves were constructed based on netDx-predicted classes of test samples. The R packages *survival* and *survminer* were used to compute Kaplan-Meier curves and *rms* was used to calculate the log-rank test for separation of survival curves. The package *survival* was also used to compute the Cox proportional hazards model of predicted poor survivors, using predicted good survivors as a reference, and to calculate the hazard ratio and associated p-value.

### Pathway networks

Pathway definitions were aggregated from HumanCyc^4^ (http://humancyc.org), NetPath^5^ (http://www.netpath.org), Reactome^6,7^ (http://www.reactome.org), NCI Curated Pathways^8^, mSigDB^9^ (http://software.broadinstitute.org/gsea/msigdb/), and Panther^10^(http://pantherdb.org/) (downloaded from http://download.baderlab.org/EM_Genesets/February_01_2018/Human/symbol/Human_AllPathways_February_01_2018_symbol.gmt)^11^. Only pathways with 10 to 500 genes were included (1,801 pathways). Pathway-level patient similarity was defined as the Pearson correlation of the expression vectors corresponding to member genes, and the network was sparsified (see next section).

### Sparsification of input networks

Edges with weights below floating-point precision were removed. The top 50 edges per node were retained (ties were ignored) to a maximum of 6,000 edges per network, following established GeneMANIA data processing procedures^12^. Where the resulting network excluded patients, the top-weighted edge for each patient was added with an edge weight at floating-point precision. The algorithm requires all patients to be in the network to allow test patients to be classified. For ovarian cancer, a less stringent sparsification method provided better performance where clinical data were included (*baserep1* model). This method applied a similarity threshold of 0.3 and included ties when keeping the 50 strongest edges per patient; in case of ties, all interactions tied with the 50th ranked interaction are retained, for a maximum of 2% of the sample size, or 600 patients^12^.

### Map of feature-selected networks

The Enrichment Map app (3.1.0RC4) in Cytoscape 3.5.1^13^ was used to generate enrichment maps^11^. A Jaccard overlap threshold of 0.05 was used to prune identical gene sets. AutoAnnotate v1.1.0 was used to cluster similar pathways using MCL clustering with default parameters.

The weighted shortest path between patient classes (a node set) was computed using Dijkstra’s method (*igraph* v1.01^14^); distance was defined as 1-similarity (or edge weight from a patient similarity network). The overall shortest path was defined as the mean pairwise shortest-path for a node set.

## Supplementary Figures

**Supplementary Figure 1.**
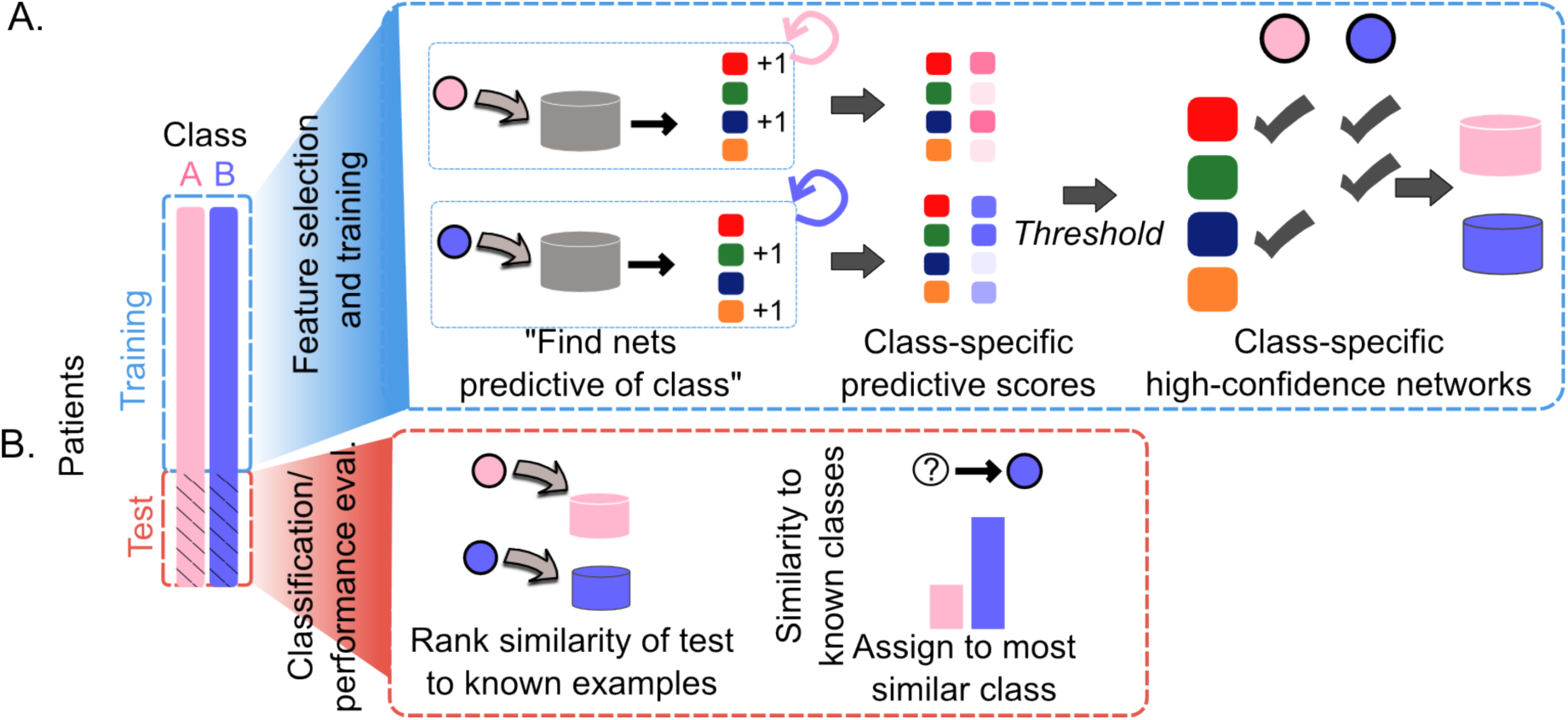
Details of the netDx feature selection and patient classification steps. A. Machine learning is used to identify networks predictive of each patient class. Data are split into training and test samples, and feature selection uses only training samples. Multiple rounds of prediction are used to score how frequently a network is predictive of a given class (e.g. high-risk). This step results in network scores, with higher values indicating networks that contribute more to prediction. These scores can be thresholded to identify a set of high-confidence networks for each class of interest (pink and blue cylinders), which represent the selected features that will be used in the final classifier. B. Test patients are ranked by similarity to known examples from the training set. For this step, only class-specific feature-selected networks are used. Patients are assigned to the class to which they have highest similarity.

**Supplementary Figure 2.**
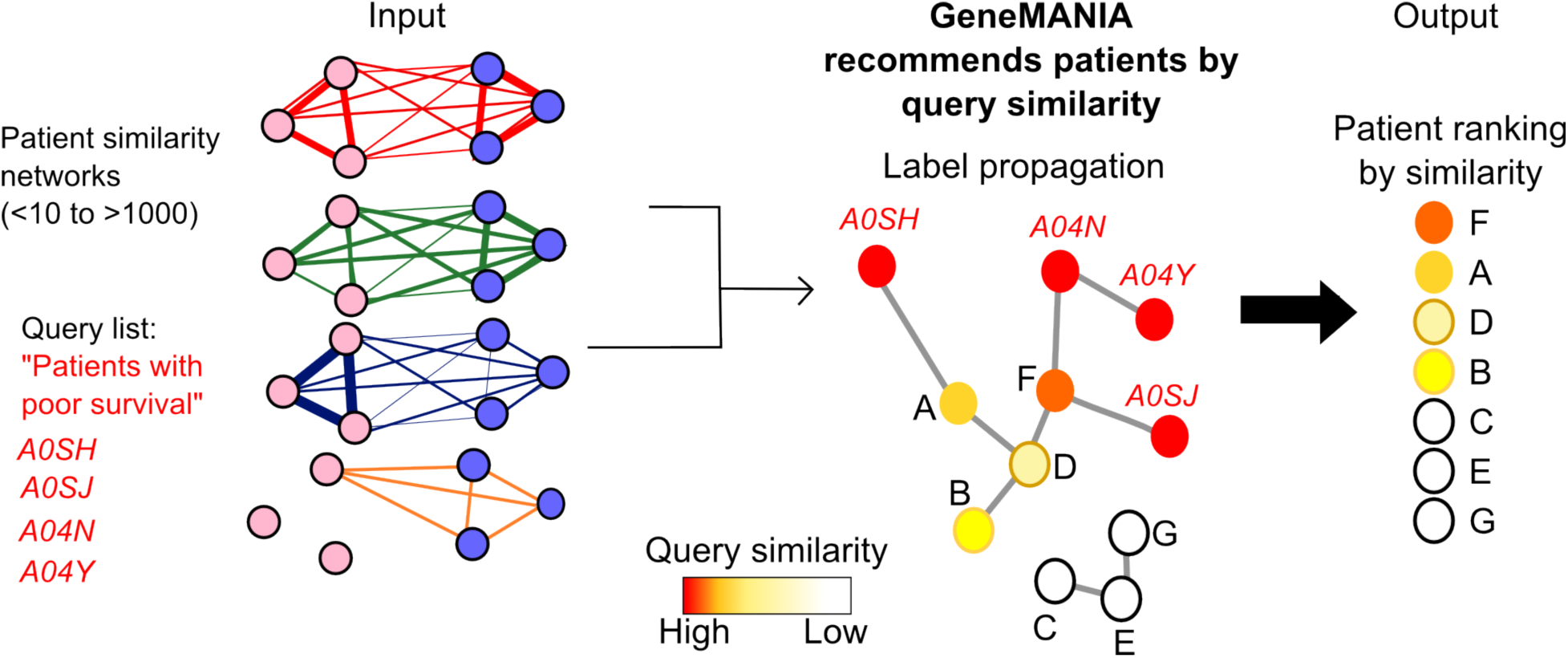
Conceptual overview of the GeneMANIA algorithm, used by netDx for network integration. GeneMANIA is a network-based recommender system that ranks all nodes by similarity to an input query (or “positive” nodes). In netDx, the nodes are patients and GeneMANIA uses the set of input patient similarity networks (left). The patient ranking is achieved by a two-step process. First, input networks are integrated into a single association network via regularized regression that maximizes connectivity between nodes with the same label and reduces connectivity to other nodes (middle); this step computes network weights corresponding to predictive value for each network. Second, label propagation is applied to the integrated network starting with the query nodes (red), thereby ranking patients from most to least similar to the query (right).

**Supplementary Figure 3.**
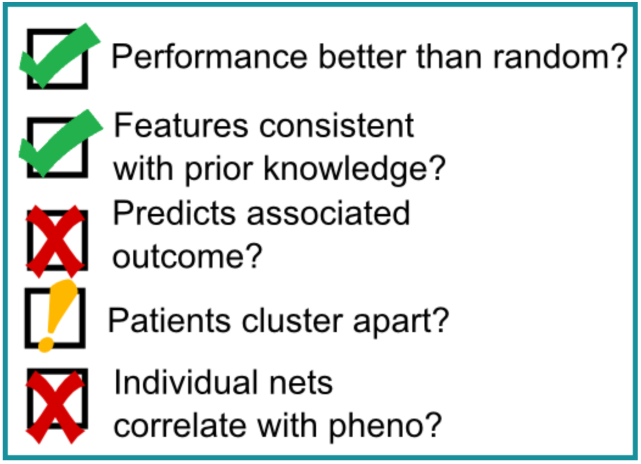
Items in a predictor checklist that could be used to compare performance of several predictors. See main text for discussion.

## Supplementary Tables

**Supplementary Table 1.**
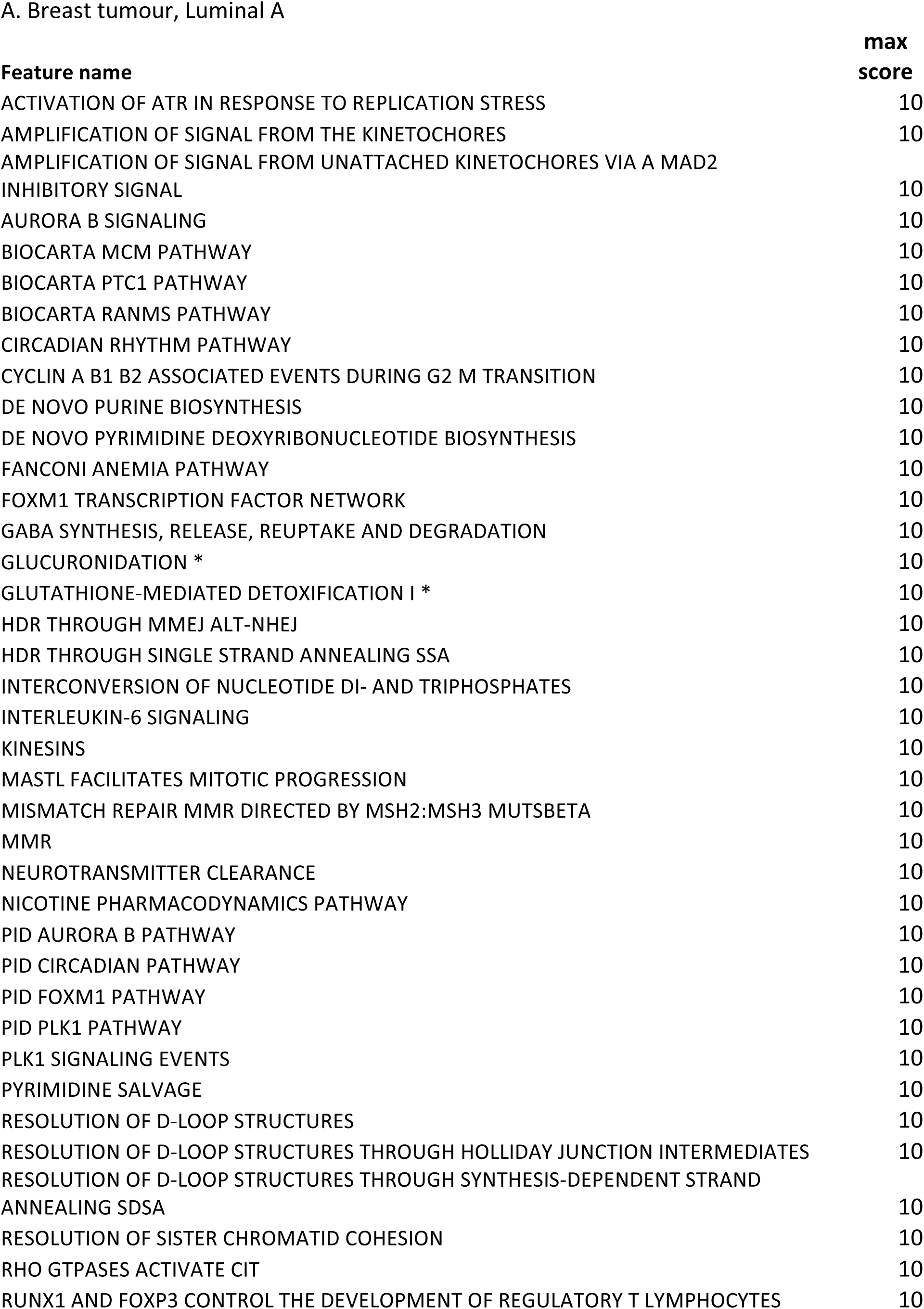
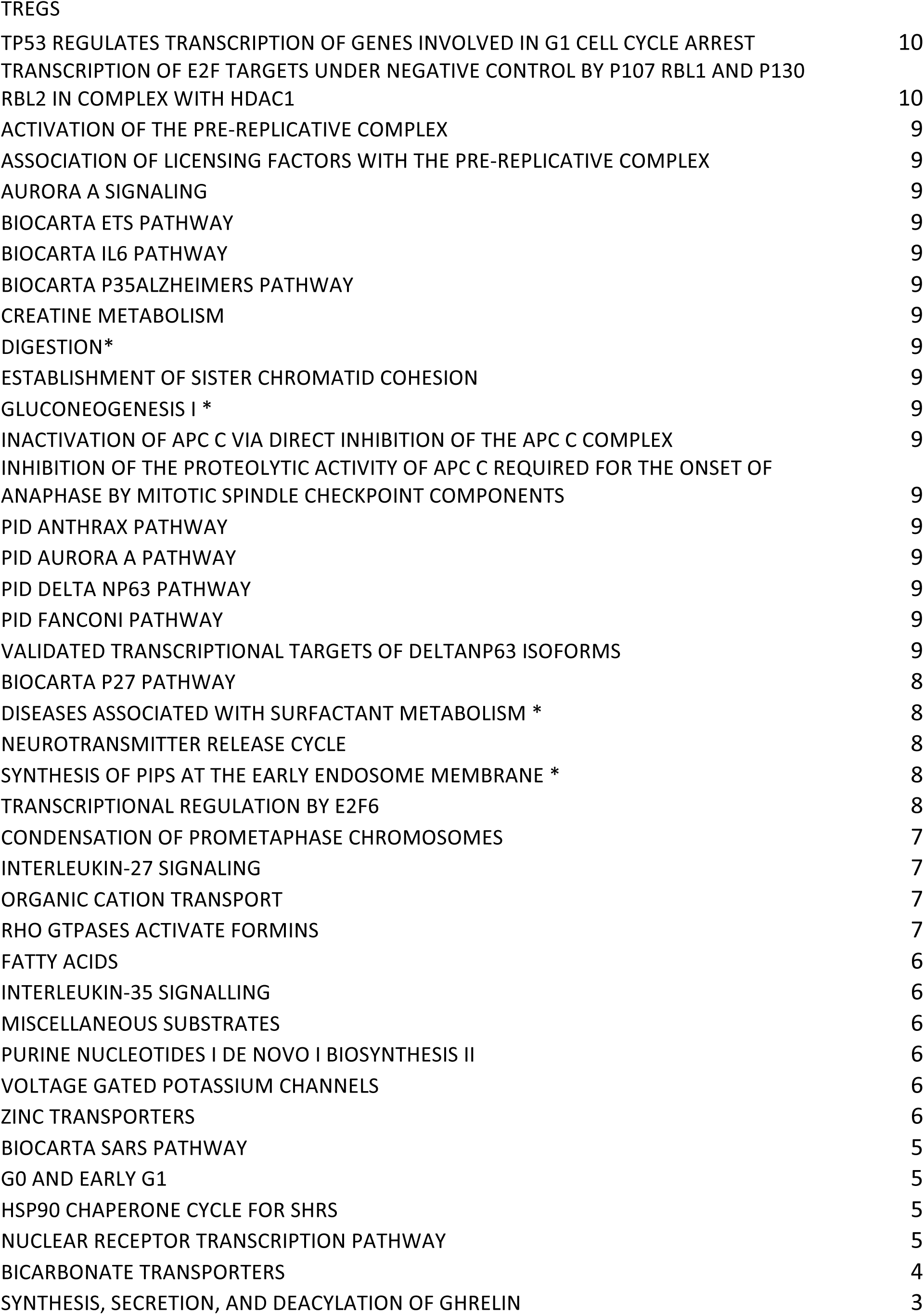
Average AUROC for netDx-predicted binarized survival prediction data for kidney, ovarian, lung and brain cancers. In each case, the value shown is the average of AUROC across 20 train/test splits.

**Supplementary Table 2.**
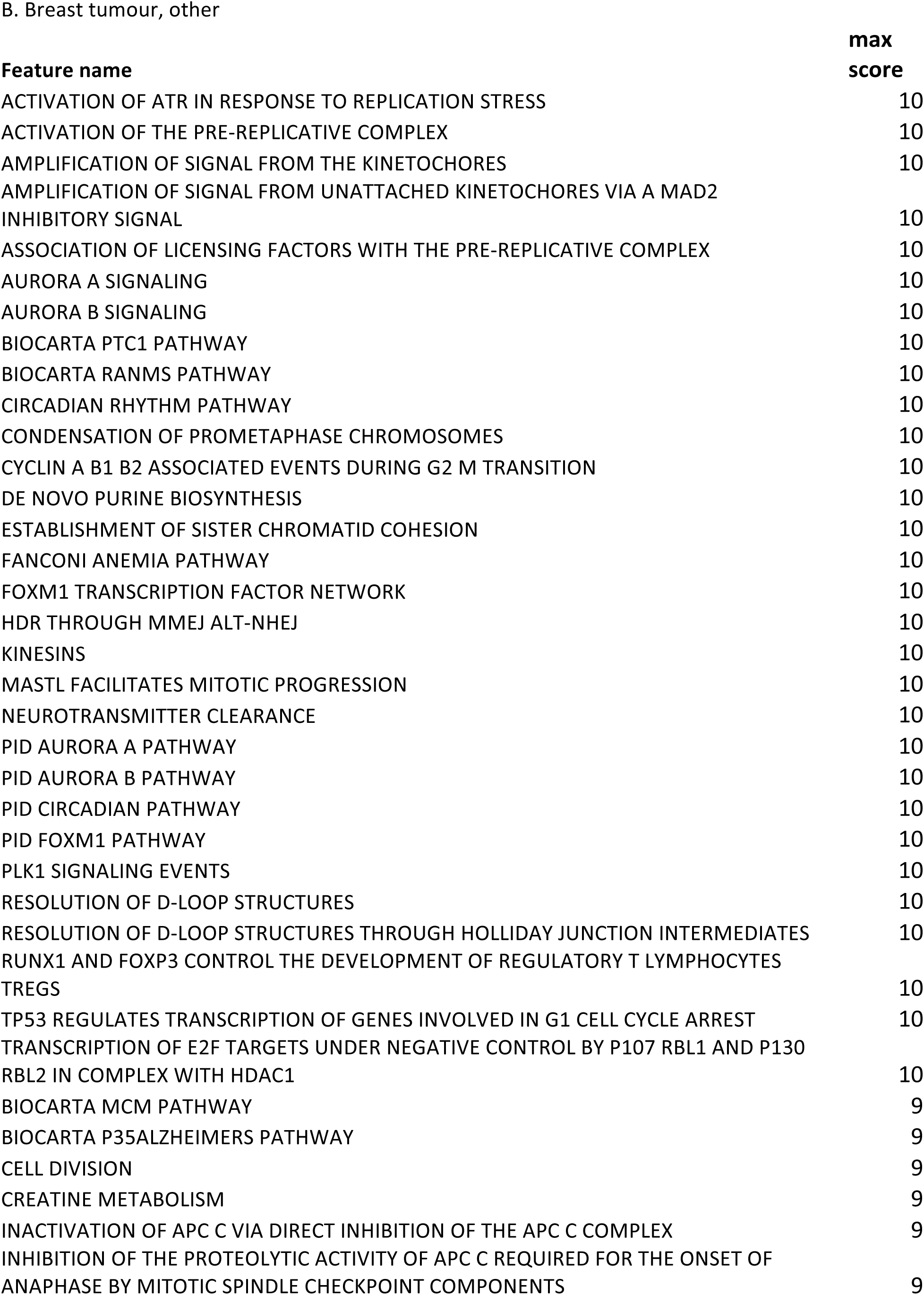
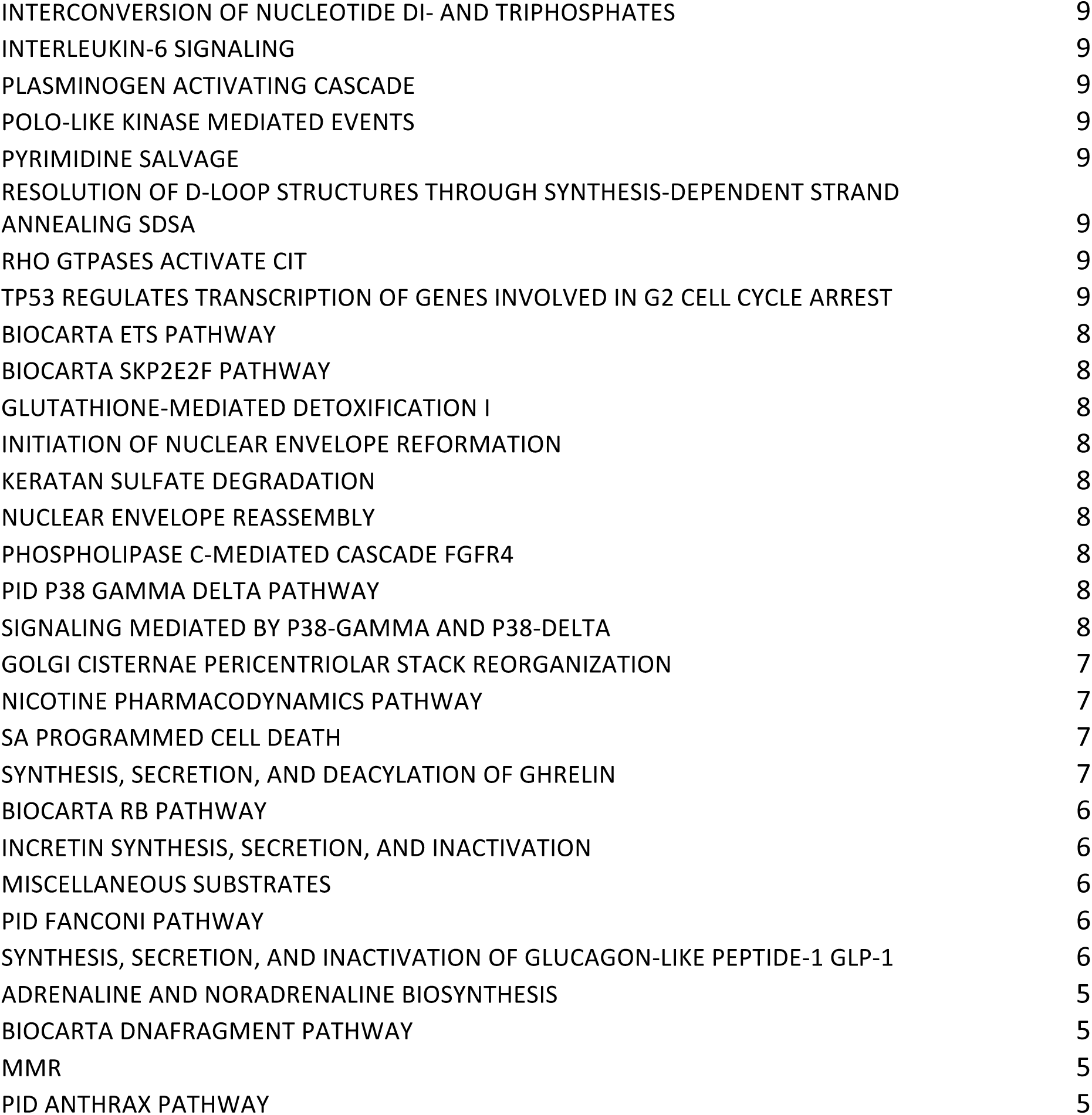
Scores for pathway-level networks for predicting Luminal A subtype of breast tumour from gene expression l. Score shown is the best achieved by a given network for over 70% of the 100 trials. Only networks scoring a max of three or more out of 10 in over 70% trials are shown here. Asterisks indicate high-scoring singleton nodes omitted from the Enrichment Map in Figure 3A.

**Supplementary Table 3.** netDx scores for pathway-level features in asthma case/control prediction. Score shown is the best achieved by a given network for over 70% of the 100 trials. Only networks scoring a max of three or more out of 10 in over 70% trials are shown here.

| Feature name | max score |
| --- | --- |
| BIOCARTA SET PATHWAY | 10 |
| BIOCARTA CTL PATHWAY | 9 |
| BIOCARTA D4GDI PATHWAY | 9 |
| NOTCH2 INTRACELLULAR DOMAIN REGULATES TRANSCRIPTION | 9 |
| SA CASPASE CASCADE | 8 |

| Feature name | max score |
| --- | --- |
| BIOCARTA CTL PATHWAY | 10 |
| BIOCARTA D4GDI PATHWAY | 10 |
| BIOCARTA SET PATHWAY | 10 |
| SA CASPASE CASCADE | 10 |
| ACTIVATION OF THE MRNA UPON BINDING OF THE CAP-BINDING COMPLEX AND EIFS, AND SUBSEQUENT BINDING TO 43S | 8 |
| BIOCARTA DNAFRAGMENT PATHWAY | 8 |
| DISEASES ASSOCIATED WITH VISUAL TRANSDUCTION | 8 |
| RETINOID CYCLE DISEASE EVENTS | 8 |

